# SNED1 modulates ECM architecture and cell proliferation via LDV-binding integrins

**DOI:** 10.64898/2026.03.16.712162

**Authors:** Dharma Pally, Leanna Leverton, Asantewaa Jones, Alexandra Naba

**Affiliations:** Department of Physiology and Biophysics, University of Illinois Chicago, Chicago, IL, 60612, U.S.A; University of Illinois Cancer Center, Chicago, IL 60612, U.S.A

**Keywords:** Cell alignment, Cell-derived matrix, Decellularization, ECM assembly, ECM fibrils

## Abstract

The extracellular matrix (ECM) is a complex scaffold of proteins that supports multicellular structures. Interactions between cells and the ECM via receptors, like integrins, govern cellular phenotypes (*e.g.*, proliferation, adhesion), but also contribute to ECM assembly. Understanding how ECM-receptor interactions regulate matrix assembly is critical to uncover how alterations of the ECM cause or accompany congenital diseases, cancer, or fibrosis.

SNED1 is a novel ECM protein with roles in development and metastasis. However, the mechanisms governing its assembly and signaling functions remain largely unknown. SNED1 contains two integrin-binding motifs, RGD and LDV, and we recently showed that its interaction with RGD-integrins mediates cell adhesion. Here, we investigated the role of SNED1/integrin interactions in SNED1 ECM assembly. While SNED1/integrin interactions were not necessary for its initial incorporation in the ECM, interaction with LDV-, but not RGD-, integrins, was required for ECM build-up and the patterning of SNED1 and the fibrillar proteins fibronectin and collagen I. Moreover, SNED1/LDV-integrin interaction promoted ECM alignment, cell alignment, and cell proliferation, processes essential to SNED1-driven neural crest cell migration during craniofacial development and breast cancer invasion.

**SUMMARY STATEMENT:** Interaction of SNED1 with LDV-binding integrins, but not RGD-binding integrins, mediates ECM remodeling and controls cytoskeletal rearrangement and cell proliferation.

## INTRODUCTION

The extracellular matrix (ECM) is a complex assembly of proteins and glycans that constitutes the architectural scaffold of multicellular structures (Hynes and Naba, 2012; Karamanos et al., 2021). Beyond physical support, the ECM provides biochemical and biomechanical cues transduced by receptors, such as the integrins, that regulate a wide range of cellular functions, including adhesion, migration, and proliferation (Chastney et al., 2025; Jones and Jones, 2023; Kanchanawong and Calderwood, 2023; Moreno-Layseca and Streuli, 2014; Pally and Naba, 2024; Saraswathibhatla et al., 2023; Yamada et al., 2019). The building of the ECM is a multi-step process that starts with the deposition of fibrillar ECM proteins, like fibronectin and collagen I, in the extracellular space (Kadler et al., 2008; Naba, 2024; Schwarzbauer and DeSimone, 2011). This initial template serves as a platform for the assembly of additional fibrillar and non-fibrillar proteins to form the mature ECM (Naba, 2024). Mechanistically, ECM build-up requires tightly regulated ECM protein-protein interactions, but also interactions between ECM proteins and their receptors, such as the integrins (Mao and Schwarzbauer, 2005; Musiime et al., 2021; Naba, 2024; Sun et al., 2025). Disruption of ECM scaffold integrity has severe consequences, ranging from congenital diseases (Lamandé and Bateman, 2020) to impaired tissue repair, cancer, and fibrosis (Bonnans et al., 2014; Lu et al., 2011). Deciphering the mechanisms underlying homeostatic ECM assembly and remodeling is thus a prerequisite to understanding how the ECM contributes to pathological processes.

We have recently identified the first functions of a novel fibrillar ECM protein, SNED1 (Sushi, Nidogen, and EGF-like domains 1), and found that it is critical for embryonic development, since *Sned1* knockout mice died shortly after birth due to craniofacial malformations (Barqué et al., 2021). We have also found that SNED1 promotes breast cancer metastasis (Naba et al., 2014), a process that, like craniofacial morphogenesis, requires structural support and signals from the ECM scaffold to guide cell adhesion, migration, and proliferation (Gallik et al., 2017; Stanger and Wahl, 2024; Walma and Yamada, 2020). Mechanistically, the SNED1 sequence contains two integrin binding motifs, RGD and LDV, and we have recently shown that SNED1 mediates cell adhesion via RGD integrins, namely α5β1 and αvβ3, but not LDV integrins, such as α4β1 (Pally et al., 2025). However, to date, we do not know how SNED1 assembles into the ECM and whether it exerts other functions beyond mediating cell adhesion.

In this short report, we investigated a possible role for integrins in SNED1 fibrillar ECM assembly. We found that mutating the two integrin-binding motifs, RGD and LDV, in SNED1, alone or in combination, did not impair initial SNED1 deposition. However, we observed that mutation of the LDV motif, but not of the RGD motif, significantly altered overall ECM architecture and build- up. In a companion study submitted with this manuscript by Leverton *et al*., we observed that SNED1 closely colocalizes with fibronectin and collagen I and requires their presence for initial fibrillar assembly in the ECM. However, we also reported that, as ECM matures *in vitro*, SNED1 fibrils separate from the fibronectin and the collagen I meshwork. Here, we found that, instead of separating from each other as observed with SNED1^WT^, SNED1^LAV^ fibers remained intertwined within the fibronectin and collagen I meshwork, suggesting a role for SNED1-LDV integrin interaction in the maturation of the ECM. Functionally, we found that SNED1^LAV^ expression perturbed ECM alignment and cell alignment and reduced cell proliferation. Altogether, our results point to a role for SNED1-LDV integrin-interactions in homeostatic ECM remodeling and signaling to orchestrate cytoskeletal rearrangement and cell proliferation.

## RESULTS AND DISCUSSION

### Establishment of a cell culture system to test the roles of SNED1 integrin-binding motifs in ECM assembly, patterning, and cellular phenotypes

To explore the possible contribution of integrins in the assembly and maturation of the architecture of the SNED1 fibrillar ECM meshwork, we engineered immortalized embryonic fibroblasts isolated from *Sned1* knockout mice (*Sned1*^KO^ iMEFs) (Barqué et al., 2021) to stably expressed GFP-tagged constructs of SNED1 in which we mutated its integrin-binding motifs: the RGD motif into RGE and the LDV motif into LAV, alone or in combination. These mutations have been extensively characterized and disrupt interactions with the eight RGD integrins (namely α5β1, α8β1, the five αv heterodimers and the platelet-specific αIIbβ3) and the eight LDV integrins (namely α4β1, α9β1, α4β7 and the leukocyte-specific β2 integrin subfamily and αEβ7), respectively (Campbell and Humphries, 2011; Hynes and Naba, 2012). Here, we confirmed that the constructs were expressed in cells (**Supplemental Figure S1A**) and that the point mutations did not impair protein secretion, as the mutant forms of SNED1 were detected in the conditioned culture media (**Supplemental Figure S1B**). Of note, we observed that SNED1^RGE^-GFP was present in higher abundance in the conditioned medium than SNED1^WT^-GFP, SNED1^LAV^-GFP, or SNED1^RGE/LAV^, whose levels were comparable (**Supplemental Figure S1B**). Last, we showed that mutated forms of SNED1 incorporated into the ECM using a sodium deoxycholate (DOC) solubility assay. In addition to being detected in the DOC-soluble protein fraction, which is enriched for intracellular proteins, SNED1^RGE^, SNED1^LAV^, and SNED1^RGE/LAV^ were detected in the DOC-insoluble protein fraction, enriched for ECM proteins, as early as 48 hours post-seeding (**Supplemental Figure S1C**). This suggests that disrupting integrin-binding sites does not impair the initial incorporation of SNED1 into the insoluble ECM. In establishing this system, we also confirmed that the fibroblasts expressed the main receptors for the RGD and LDV motifs, namely α5β1 and α4β1, and that overexpression of the mutated forms of SNED1 did not affect the abundance of the different integrin chains (**Supplemental Figure S1D**).

### Disruption of the RGD and LDV integrin-binding motifs in SNED1 does not impair the early step of ECM assembly

Using this experimental system, we first examined whether mutations in the integrin-binding sites of SNED1 affected its incorporation into the ECM scaffold. Using confocal immunofluorescence microscopy, we monitored the assembly and organization of SNED1 and two major fibrillar ECM proteins, fibronectin and collagen I, in the ECM produced by cells for 3 days in culture. We found that neither the disruption of the RGD motif nor the disruption of the LDV motif altered the ability of SNED1 to incorporate in the ECM (**Figure 1A and 1C**). We also found that expression of the mutant forms of SNED1 did not alter the patterning of fibronectin (**Figure 1A**) or collagen I (**Supplemental Figure S2A**). Expression of integrin-binding mutant forms of SNED1 did not alter the colocalization of SNED1 with fibronectin (**Figures 1B-1D**) or collagen I (**Supplemental Figures S2B-S2D**) that we reported in the companion study by Leverton *et al*.

**Figure 1.**
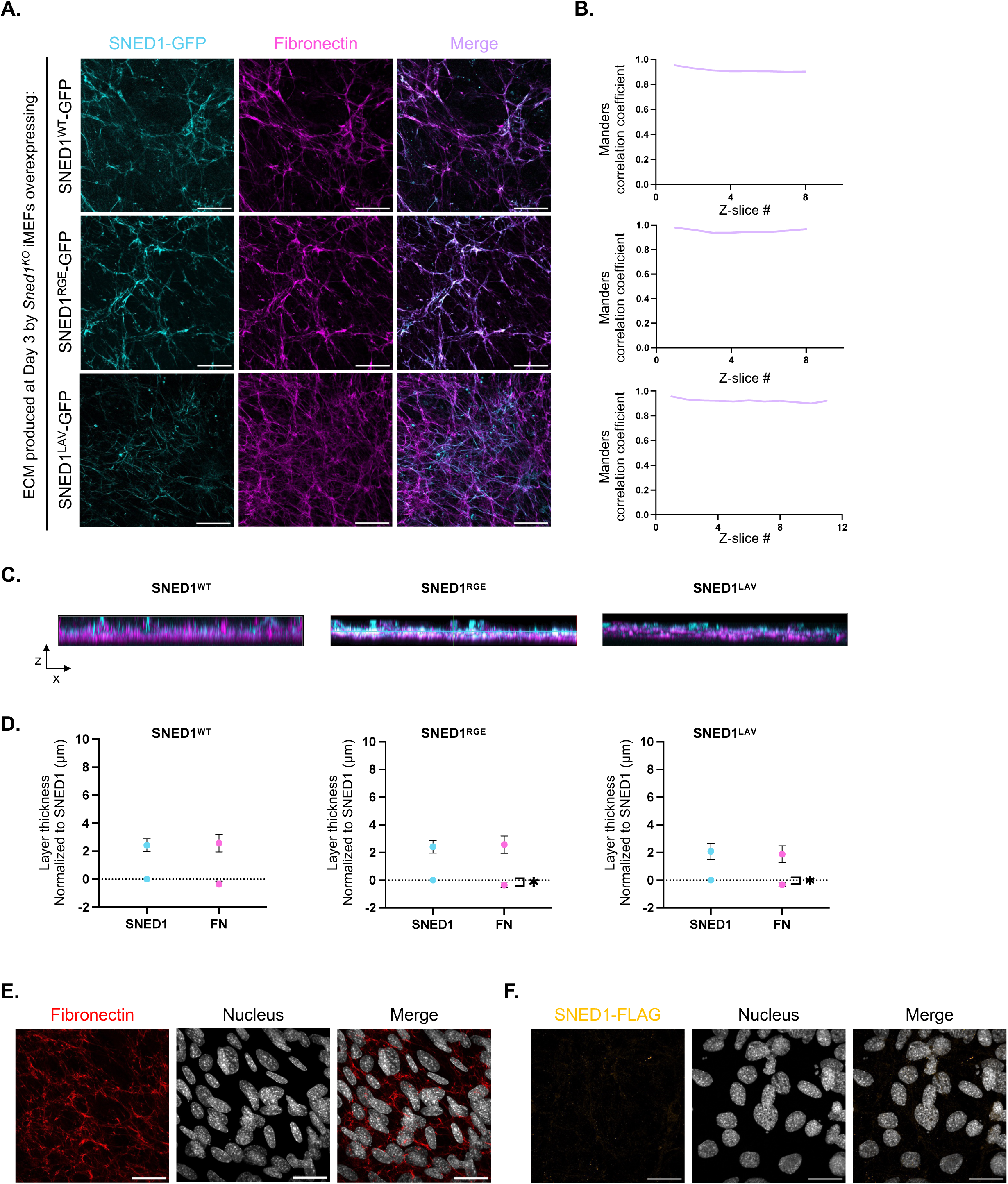
Disruption of the RGD or LDV integrin-binding motifs in SNED1 does not affect the early stages of ECM assembly or patterning of SNED1. **A.** XY orthogonal maximum projections show SNED1 (cyan) and fibronectin (magenta) fibers in ECMs produced by *Sned1*^KO^ iMEFs overexpressing SNED1^WT^-GFP (*top panels*), SNED1^RGE^-GFP (*middle panels*), or SNED1^LAV^-GFP (*bottom panels*) and decellularized 3 days post-seeding. Merge panels show overlap between SNED1 and fibronectin signals. Scale bar: 30 µm. Images are representative of at least three independent biological replicates. **B.** Representative line graphs show Manders overlap coefficient between SNED1 and fibronectin for each Z-slice where in-focus signal was detected in the ECM produced by *Sned1*^KO^ iMEFs overexpressing SNED1^WT^-GFP (*top panel*), SNED1^RGE^-GFP (*middle panel*) or SNED1^LAV^-GFP (*bottom panel*) and decellularized 3 days post-seeding. **C.** XZ orthogonal maximum projections show fibronectin and SNED1-GFP signal in the ECMs produced by *Sned1*^KO^ iMEFs overexpressing SNED1^WT^-GFP (*left panel*), SNED1^RGE^-GFP (*middle panel*), or SNED1^LAV^-GFP (*right panel*), decellularized 3 days post cell-seeding. Images are representative of three independent biological replicates. **D.** Dot plots depict the overlap between the SNED1 and fibronectin (FN) signals in ECMs produced by *Sned1*^KO^ iMEFs overexpressing SNED1^WT^-GFP (*left panel*), SNED1^RGE^-GFP (*middle panel*) or SNED1^LAV^-GFP (*right panel*) and decellularized 3 days post-seeding. Lower and higher data points represent the beginning and end of in-focus signal detected in the ECM. Data points are normalized to the Z-slice value where SNED1’s signal is detected. Data is represented as mean ± standard deviation. Unpaired *t*-test with Welch’s correction was performed to test statistical significance. *p<0.05. **E.** XY orthogonal maximum projections show fibril assembly of exogenously added fibronectin (red) by *Sned1*^KO^ iMEFs overexpressing GFP alone (nucleus staining; grey). Merge panels show cells and fibronectin fibrils signals. Images are representative of two independent biological replicates. Scale bar: 30 µm. **F.** XY orthogonal maximum projections of *Sned1*^KO^ iMEFs overexpressing GFP alone (nucleus staining; grey) incubated in presence of purified SNED1^WT^-FLAG (orange). Images are representative of two independent biological replicates. Scale bar: 30 µm.

To more definitively rule out the involvement of integrins in the early step of SNED1 assembly, we asked whether cells were capable of assembling SNED1 fibers if incubated in presence of exogenous SNED1. To do so, we supplemented the culture medium of *Sned1*^KO^ iMEFs overexpressing GFP alone with exogenous fibronectin, a protein known to require interaction with integrins to initiate its fibrillar assembly, or SNED1^WT^-FLAG. After 3 days in culture, unlike fibronectin (**Figure 1E**), we did not observe SNED1 fibrils (**Figure 1F**).

Together with the observations reported in the companion paper by Leverton *et al*., our findings demonstrate that the initial step of SNED1 incorporation into the ECM is integrin-independent and instead relies on the presence of fibrillar ECM proteins, fibronectin and collagen I. Interestingly, this represents a distinct mechanism from integrin-mediated initial ECM assembly of other ECM proteins, such as that of fibronectin, where fibronectin fibril assembly is initiated by the interaction of α5β1 integrins with its RDG motif before involving ECM protein-ECM protein interactions (Mao and Schwarzbauer, 2005; Singh et al., 2010).

### Disruption of the LDV but not the RGD integrin-binding motif in SNED1 impairs ECM remodeling and SNED1 distribution within mature ECMs

As discussed in the companion paper to this study, we found that after early assembly, ECM produced by immortalized mouse embryonic fibroblasts undergoes remodeling and that the meshwork of SNED1^WT^ fibers, which initially overlaps with fibronectin and collagen I (**Figure 1**), separates from the meshwork containing collagen I and fibronectin (**Figures 2A-2D**).

**Figure 2.**
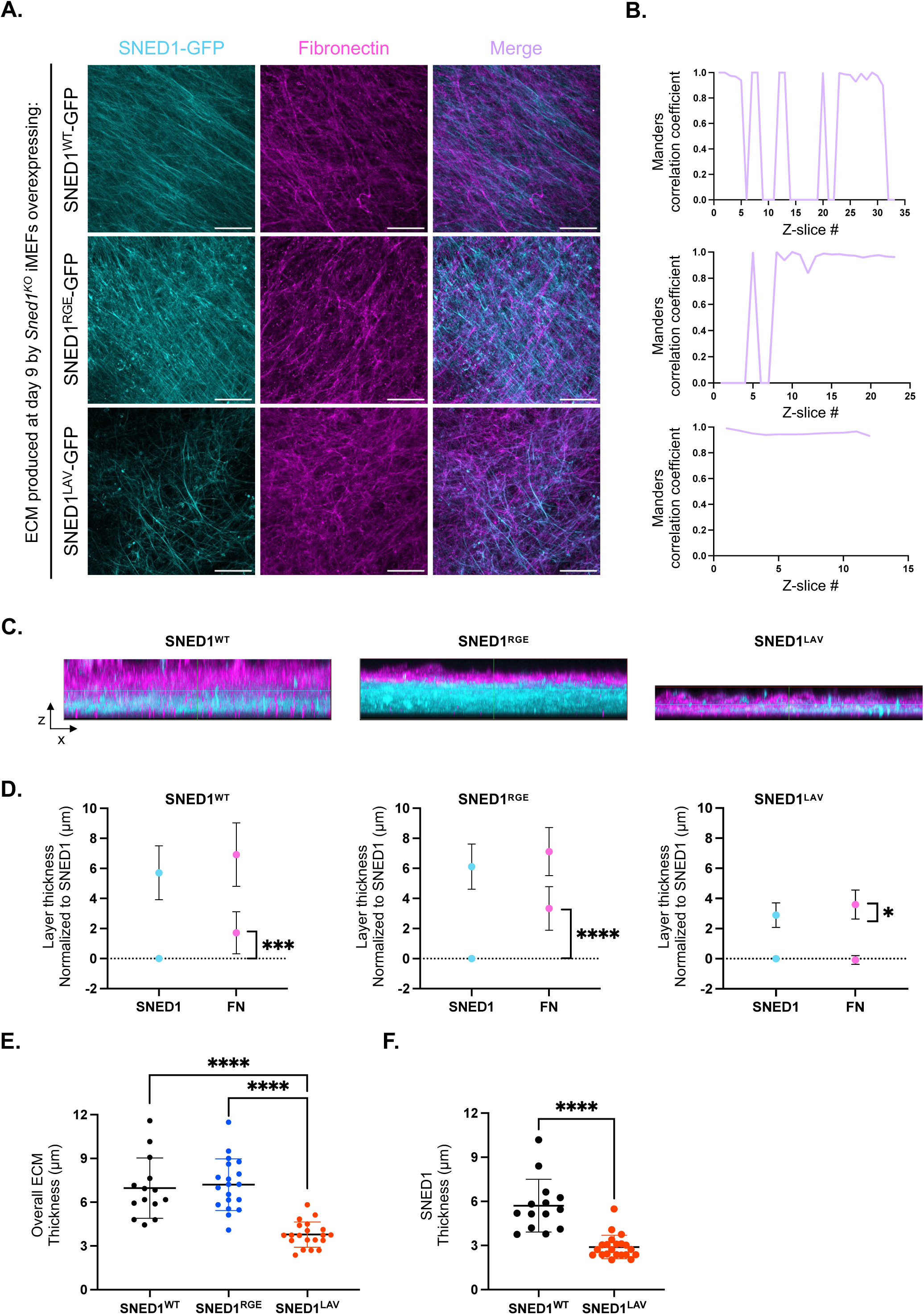
Disruption of the LDV but not the RGD integrin-binding motif in SNED1 impairs ECM remodeling. **A.** XY orthogonal maximum projections show SNED1 (cyan) and fibronectin (magenta) fibers in the ECM produced by *Sned1*^KO^ iMEFs overexpressing SNED1^WT^-GFP (*top panels*), SNED1^RGE^- GFP (*middle panels*), or SNED1^LAV^-GFP (*bottom panels*) and decellularized 9 days post-seeding. Merge panels show overlap between SNED1 and fibronectin signals. Scale bar: 30 µm. Images are representative of at least three independent biological replicates. **B.** Representative line graphs show Manders overlap coefficient between SNED1 and fibronectin for each Z-slice where in-focus signal was detected in the ECM produced by *Sned1*^KO^ iMEFs overexpressing SNED1^WT^-GFP (*top panel*), SNED1^RGE^-GFP (*middle panel*), or SNED1^LAV^-GFP (*bottom panel*), decellularized 9 days post-seeding. **C.** XZ orthogonal maximum projections show fibronectin (magenta) and SNED1-GFP (yellow) signals in the ECMs produced by *Sned1*^KO^ iMEFs overexpressing SNED1^WT^-GFP (*left panel*), SNED1^RGE^-GFP (*middle panel*), or SNED1^LAV^-GFP (*right panel*) decellularized at 9 days post cell-seeding. Images are representative of three independent biological replicates. **D.** Dot plots depict the overlap between SNED1 and fibronectin (FN) signals in ECMs produced by *Sned1*^KO^ iMEFs overexpressing SNED1^WT^-GFP (*left panel*), SNED1^RGE^-GFP (*middle panel*), or SNED1^LAV^-GFP (*right panel*) decellularized at 9 days post-seeding. Lower and higher data points represent the beginning and end of in-focus signal detected in the ECM. Data points are normalized to the Z-slice value where SNED1’s signal is detected. Data is represented as mean ± standard deviation. Unpaired *t*-test with Welch’s correction was performed to test statistical significance. *p<0.05, ***p<0.001, ****p<0.0001. **E.** Dot plot depicts the overall thickness of ECMs produced by *Sned1*^KO^ iMEFs overexpressing SNED1^WT^-GFP, SNED1^RGE^-GFP, or SNED1^LAV^-GFP decellularized at 9 days post-seeding. Data is represented as mean ± standard deviation from three biological replicates, with at least 4 imaged fields per replicate and timepoint. Unpaired *t*-test with Welch’s correction was performed to test statistical significance. ****p<0.0001. **F.** Dot plot depicts SNED1 thickness in the ECMs produced by *Sned1*^KO^ iMEFs overexpressing SNED1^WT^-GFP or SNED1^LAV^-GFP decellularized at 9 days post-seeding. Data is represented as mean ± standard deviation from three biological replicates, with at least 4 imaged fields per replicate and timepoint. Unpaired *t*-test with Welch’s correction was performed to test statistical significance. ****p<0.0001.

Similarly to what we observed upon overexpression of SNED1^WT^, we observed that in mature ECMs produced by cells overexpressing SNED1^RGE^ for 9 days in culture, the meshwork of SNED1^RGE^ fibers separates from the fibronectin (**Figures 2A-2D**) and collagen meshworks (**Supplemental Figures S3A-S3D**). Similarly to SNED1^WT^, we observed a decrease in the overlap between the SNED1^RGE^-GFP and fibronectin (**Figures 2A and 2B**) or collagen I (**Supplemental Figures S3A and S3B**). Assessment of the distribution of the proteins through the thickness of the ECM using XZ orthogonal maximum intensity projection revealed that in ECMs produced by cells expressing SNED1^WT^ or SNED1^RGE^, the most basal portion of the ECM is devoid of fibronectin; conversely the upper portion of the ECM (∼1.5 µm) exclusively contains fibronectin and is devoid of SNED1 (**Figures 2C and 2D**).

In stark contrast, ECM remodeling and partitioning of different ECM layers did not occur upon expression of SNED1^LAV^. The quantitative analysis of the spatial overlap, in the XY (**Figure 2B**) and in the XZ (**Figure 2D**) dimensions, showed that SNED1^LAV^ and fibronectin colocalized and occupied the same portion of the ECM (**Figures 2B and 2D**, respectively), although the fibronectin extended a bit further (∼0.7 µm) than the SNED1 staining apically upper mean of fibronectin slightly but statistically significantly shifted upward (**Figure 2D**). Similar observations were made with collagen I (**Supplemental Figures S3A-S3D**). In addition, we observed that the expression of SNED1^LAV^ prevented ECM build-up and remodeling and resulted in overall thinner ECM (∼3.8 µm; **Figure 2E**) and thinner SNED1 layer (∼2.9 µm; **Figure 2F**), suggesting that this mutant acts as a dominant negative protein on ECM remodeling and maturation. Of note, the combined disruption of the two integrin-binding sites in SNED1 (SNED1^RGE/LAV^) phenocopied the SNED1LAV phenotype (**Supplemental Figures S3E and S3F**).

In light of our previous findings that the RGD but not the LDV motif in SNED1 mediates cell adhesion (Pally et al., 2025), we propose that SNED1-integrin interaction sites exert different roles. Specifically, our results point to a role for the LDV motif of SNED1, and hence its possible interaction with α4β1 integrin, in remodeling the ECM meshwork and establishing the partitioning of different ECM layers. Interestingly, Sechler *et al*., have previously found that, in absence of an RGD motif, fibronectin fibril formation can still occur but through an alternative mechanism engaging α4β1 integrin interaction with fibronectin’s LDV motif (Sechler et al., 2000). However, in SNED1, the function of the LDV motif in ECM remodeling does not appear to be redundant with that of the RGD motif.

### Mutation of the LDV integrin-binding motif affects ECM thickness and fiber alignment

A closer examination of the pattern of distribution of SNED1 in a mature (9-day old) ECM revealed that, in contrast to highly aligned SNED1^WT^, SNED1^LAV^ fibers are more disorganized and do not follow a specific alignment pattern (**Figure 3A**). Quantification of SNED1 fiber orientation showed that SNED1^WT^ fibers displayed a bimodal distribution with two distinct peaks, while SNED1^LAV^ fibers showed a wider distribution of angles with no distinct peaks (**Figure 3B**). We also found that the density of SNED1-positive staining was significantly reduced in ECM produced by cells expressing SNED1^LAV^ as compared to the ECM produced by cells expressing SNED1^WT^ (**Figure 3C**). Importantly, and as previously shown, these changes cannot be attributed to differential protein abundance, since SNED1^WT^ and SNED1^LAV^ are produced and secreted in the same proportion by cells (**Supplemental Figure S1A-C**). These cells also produce fibronectin in the same proportion (**Supplemental Figure S1E**).

**Figure 3.**
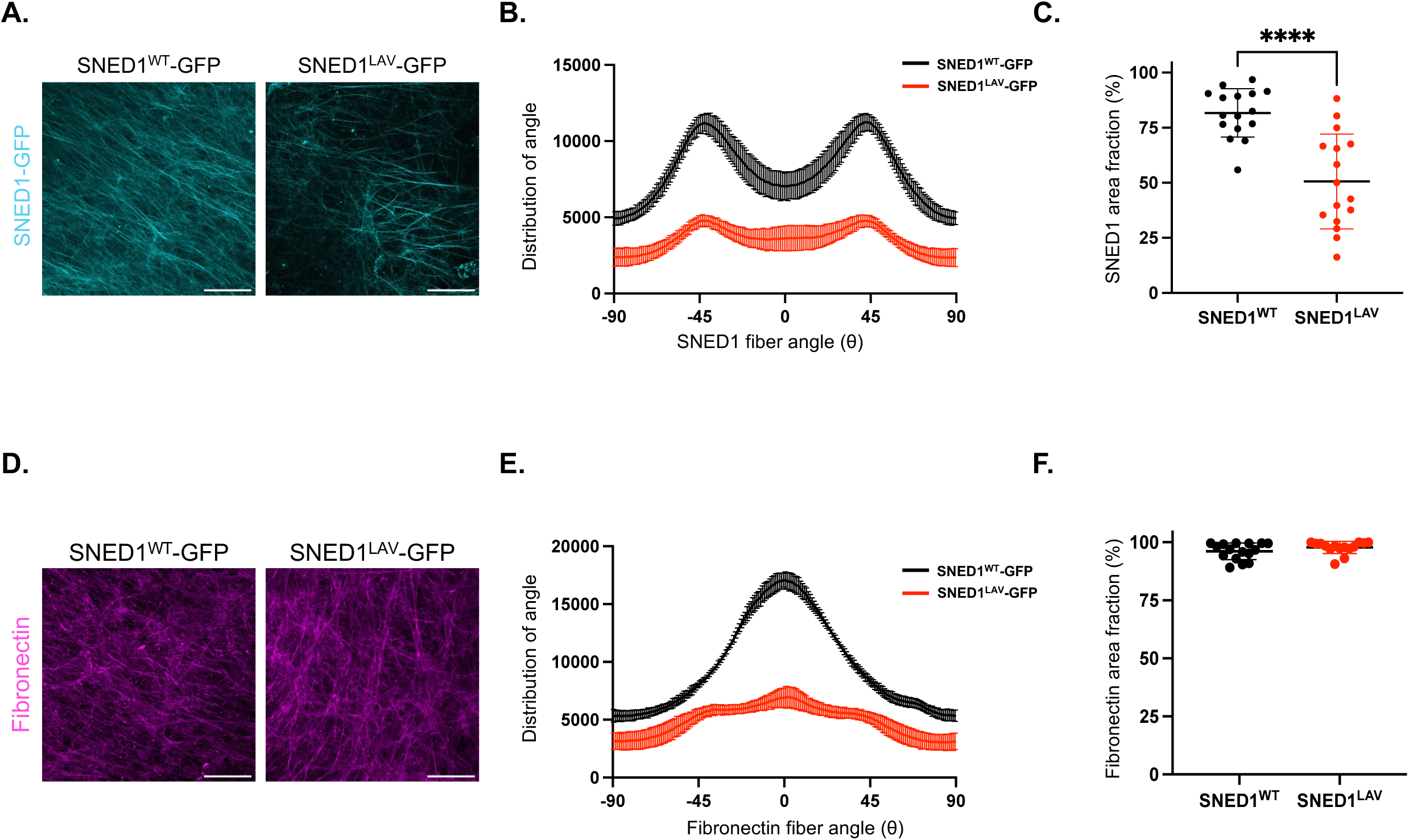
Mutation of the LDV-integrin binding motif in SNED1 results in a thinner and less aligned ECM. **A.** XY orthogonal maximum intensity projections show SNED1 fibers (cyan) in ECMs produced by *Sned1*^KO^ iMEFs overexpressing SNED1^WT^-GFP (*left panel*) or SNED1^LAV^-GFP (*right panel*) and decellularized at 9 days post-seeding. Scale bar: 30 μm. Images are representative of three independent biological replicates. **B.** Histogram depicts SNED1 fiber alignment in ECMs produced by *Sned1*^KO^ iMEFs overexpressing SNED1^WT^-GFP (black) or SNED1^LAV^-GFP (red). Data is represented as the mean ± standard deviation from three independent biological replicates with at least four non- overlapping images from each biological replicate. **C.** Dot plot shows area fraction (density) of SNED1 fibers in decellularized ECMs produced by *Sned1*^KO^ iMEFs overexpressing either SNED1^WT^-GFP (black) or SNED1^LAV^-GFP (red). Data is represented as mean ± standard deviation from three biological replicates, with at least 4 imaged fields per replicate. Unpaired *t*-test with Welch’s correction was performed to test statistical significance. ****p<0.0001. **D.** XY orthogonal maximum intensity projections show fibronectin fibers (magenta) in ECMs produced by *Sned1*^KO^ iMEFs overexpressing SNED1^WT^-GFP (*left panel*) or SNED1^LAV^-GFP (*right panel*) and decellularized at 9 days post-seeding. Scale bar: 30 μm. Images are representative of three independent biological replicates. **E.** Histogram depicts fibronectin fiber alignment in ECMs produced by *Sned1*^KO^ iMEFs overexpressing SNED1^WT^-GFP (black) or SNED1^LAV^-GFP (red). Data is represented as the mean ± standard deviation from three independent biological replicates with at least four non- overlapping images from each replicate. **F.** Dot plot shows area fraction (density) of fibronectin signal in decellularized ECMs produced by *Sned1*^KO^ iMEFs overexpressing either SNED1^WT^-GFP (black) or SNED1^LAV^-GFP (red). Data is represented as mean ± standard deviation from three biological replicates, with at least 4 imaged fields per biological replicate. Unpaired *t*-test with Welch’s correction was performed to test statistical significance (ns).

The mutation of the LDV integrin-binding motif not only affected the architecture of the SNED1 fibers but also impaired the alignment of fibronectin fibers (**Figures 3D and 3E**), again pointing to a dominant negative role for this mutant proteoform. Of note, the density of fibronectin-positive staining was not affected in ECM produced by cells expressing SNED1^LAV^ in comparison to the ECM produced by cells expressing SNED1^WT^ (**Figure 3F**). Altogether, our observations demonstrate a role for SNED1’s LDV integrin-binding motif not only in the patterning of SNED1, but also the regulation of overall ECM build-up and patterning.

The precise molecular mechanisms linking the disruption of SNED1-LDV integrin interactions with altered ECM patterning remain to be elucidated. Since several other proteins, including fibronectin but also non-ECM proteins like transmembrane cell adhesion molecules contain LDV motifs and can engage LDV integrins (Campbell and Humphries, 2011; Clements et al., 1994; LaFoya et al., 2018; Makarem and Humphries, 1991), it is possible that if SNED1 cannot engage LDV integrins, these integrins can engage with other ligands and, as a result, trigger the phenotypes observed (*e.g.*, inhibition of ECM build-up and non-partitioning of SNED1 layer from fibronectin and collagen I layer). At this stage, this hypothesis is very speculative, and additional studies will be needed to determine the mechanisms of interactions and competitions between LDV integrins and its ligands. In addition, the loss of ECM fiber alignment is often correlated with a defective mechanotransduction, the transmission of extracellular physical and mechanical signals into intracellular biochemical signals. This has been most extensively characterized for fibronectin fiber alignment through RGD-dependent integrins (Geiger et al., 2009; Lemmon et al., 2009). Interestingly, the LDV motif, a recognition site for several integrins, including α4β1 expressed by fibroblasts but also mammary tumor cells and neural crest cells (Pally et al., 2025), was found to facilitate fibronectin assembly in the absence of the RGD motif, largely by transmitting intracellular contractile forces largely through cortical actin (Sechler et al., 2000). However, to the best of our knowledge, there is no prior evidence linking the LDV motif in other ECM proteins to fiber remodeling, making SNED1 possibly unique in this sense. Furthermore, how a possible interaction with LDV integrins can cause the separation of SNED1 fibrils from the fibronectin- and collagen I-containing ECM meshwork, also remains to be elucidated. A first step toward this goal would be to demonstrate whether SNED1 indeed directly interacts with LDV integrins, in particular α4β1, and determine whether SNED1 and fibronectin can compete for the same receptors and if so, according to what binding parameters.

### ECM containing SNED1^LAV^ alters cell alignment

Integrins, by connecting cells to the surrounding ECM, serve as adhesion receptors and promote cell spreading and alignment along ECM fibers. We have previously reported that SNED1 mediates the adhesion of mammary epithelial cells and neural crest cells through its RGD motif and via α5β1 and the αvβ3 integrins, but that the LDV motif in SNED1 and α4β1 were not required for the adhesion of these cell types (Pally et al., 2025). Since we found that mutation of the LDV integrin-binding site in SNED1 led to altered ECM alignment, we next sought to determine the impact of this mutation on cellular phenotypes.

In agreement with our original findings on mammary tumor cells and neural crest cells, we found that MEFs seeded on SNED1^LAV^-coated surface could adhere to the same extent as MEFs seeded on SNED1^WT^-coated surface (**Figure 4A**). Similarly, MEFs adhered to the same extent to decellularized ECM produced by cells overexpressing SNED1^WT^ than to the ECM produced by cells overexpressing SNED1^LAV^ (**Figure 4B**). These observations rule out a role for SNED1-LDV integrin interaction, in cell adhesion. However, we observed that cells seeded on ECMs containing SNED1^LAV^ tended to spread less than those seeded on ECMs containing SNED1^WT^. Yet, quantification of different morphometric parameters, such as cell perimeter, aspect ratio, circularity, and roundness, did not meet statistical significance (**Supplemental Figure S4**).

**Figure 4.**
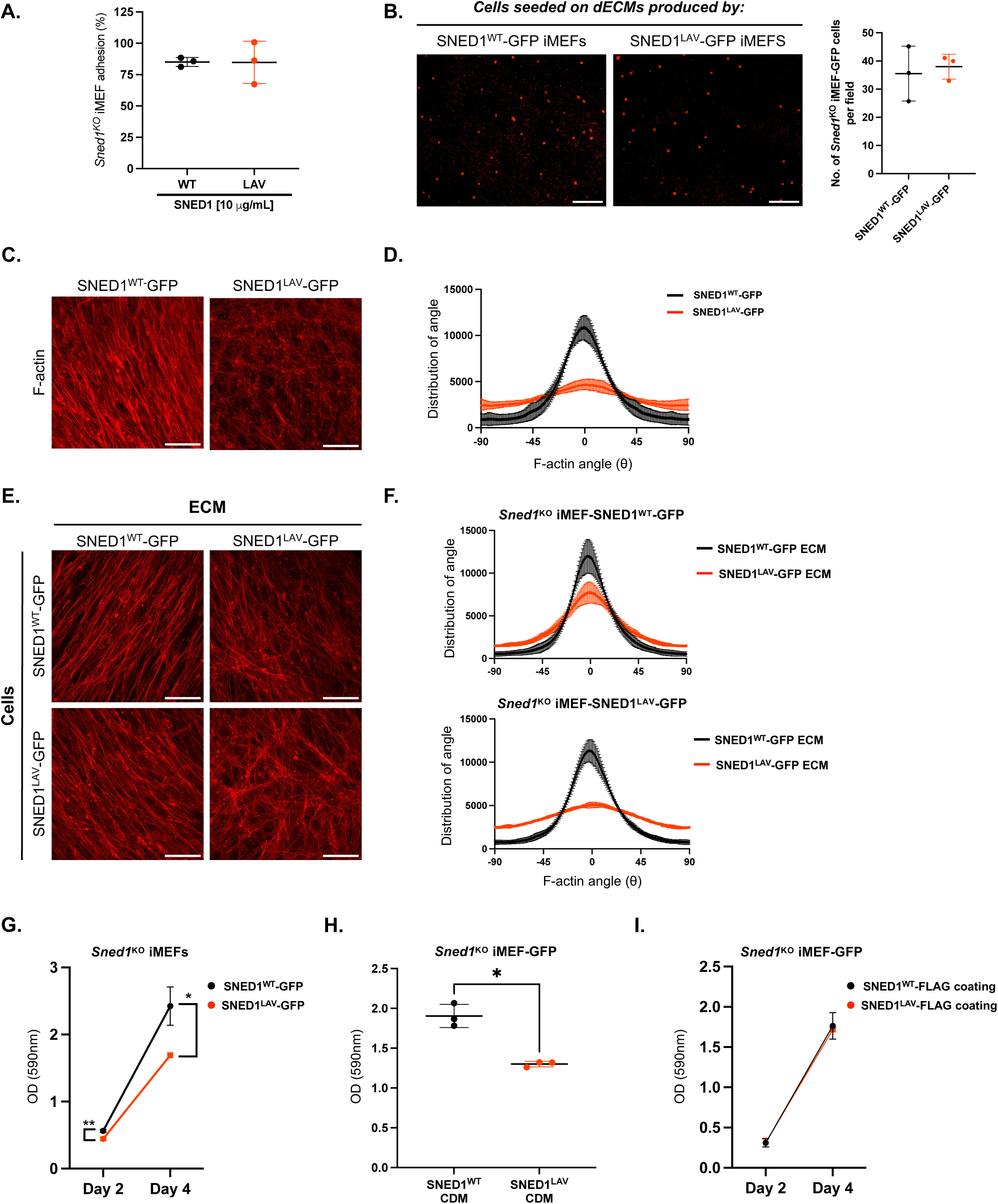
Mutation of the LDV integrin-binding motif in SNED1 impairs cell alignment and cell proliferation. **A.** Dot plot shows adhesion of *Sned1*^KO^ iMEFs on SNED1^WT^-His- (black) or SNED1^LAV^-His- (red) coated substrates. Data is represented as mean ± standard deviation from three biological replicates. Unpaired *t*-test with Welch’s correction did not show any statistical significance. **B. *Left panels:*** Microscopy images show cells (red) that have adhered after 30 min to ECMs produced by cells overexpressing SNED1^WT^-GFP (*left panel*) or SNED1^LAV^-GFP (*right panel*) and decellularized at 9 days post-seeding. Images representative of three biological replicates. Scale bar: 200 µm. ***Right panel:*** Dot plot depicts the number of adherent cells per field on decellularized ECM containing SNED1^WT^ (black) or SNED1^LAV^ (red). Data is represented as mean ± standard deviation from three biological replicates, with at least 10 fields imaged per coverslip and at least two coverslips per replicate. Unpaired *t*-test with Welch’s correction did not show any statistical significance. **C.** XY orthogonal maximum intensity projections show F-actin staining (red) in *Sned1^KO^* iMEFs overexpressing SNED1^WT^-GFP (*left panel*) or SNED1^LAV^-GFP (*right panel)* at 9 days post seeding. Scale bar: 50 μm. Images are representative of three independent biological replicates. **D.** Histogram depicts F-actin alignment in *Sned1*^KO^ iMEFs overexpressing SNED1^WT^-GFP or SNED1^LAV^-GFP at 9 days post seeding. Data is represented as the mean ± standard deviation from three independent biological replicates with at least four non-overlapping images from each replicate. **E.** XY orthogonal maximum intensity projections show F-actin staining (red) of *Sned1*^KO^ iMEFs overexpressing SNED1^WT^-GFP (*top panel*) or SNED1^LAV^-GFP (*bottom panel*) when cultured for 2 days on ECMs produced by *Sned1*^KO^ iMEFs overexpressing SNED1^WT^-GFP or SNED1^LAV^-GFP and decellularized at 9 days post-seeding. Images are representative of three independent biological replicates. Scale bar: 50 μm. **F.** Histogram show F-actin alignment in *Sned1*^KO^ iMEFs overexpressing SNED1^WT^-GFP (*top panel*) or SNED1^LAV^-GFP (*bottom panel*) when cultured on decellularized ECMs containing SNED1^WT^-GFP or SNED1^LAV^-GFP. Data is represented as mean ± standard deviation from three independent biological replicates with at least four non-overlapping images from each replicate. **G.** Line graph shows proliferation of *Sned1*^KO^ iMEFs overexpressing SNED1^WT^-GFP (black) or SNED1^LAV^-GFP (red) at 2 and 4 days post-seeding. Data is represented as mean ± standard deviation from three biological replicates. Unpaired *t*-test with Welch’s correction was performed to test statistical significance. *p<0.05. **H.** Dot plot shows proliferation of *Sned1*^KO^ iMEFs overexpressing GFP at 4 days when cultured on ECMs produced by *Sned1*^KO^ iMEFs overexpressing SNED1^WT^-GFP (black) or SNED1^LAV^-GFP (red) and decellularized at 6 days post-seeding. Data is represented as mean ± standard deviation from three biological replicates. Unpaired *t*-test with Welch’s correction was performed to test statistical significance. *p<0.05. **I.** Line graph shows proliferation of *Sned1*^KO^ iMEFs overexpressing GFP at 2 and 4 days post seeding on purified SNED1^WT^-FLAG (black) or SNED1^LAV^-FLAG (red) substrate. Data is represented as mean ± standard deviation from three biological replicates. Unpaired *t*-test with Welch’s correction did not show any statistical significance.

Since we showed that mutation of the SNED1 LDV motif altered ECM alignment and patterning, we next asked whether cells followed the same pattern. We thus quantified the orientation of actin filaments in cells expressing SNED1^WT^ or SNED1^LAV^ after 9 days in culture and found that cells expressing SNED1^LAV^ were much more disorganized than cells expressing SNED1^WT^ (**Figures 4C and 4D**). This phenotype mirrors the data reported in **Figure 3** on SNED1 and fibronectin fiber alignment patterns.

To determine whether ECM architecture can dictate cell alignment, we performed a cross-seeding experiment whereby we cultured cells overexpressing SNED1^WT^ or SNED1^LAV^ on decellularized ECM produced by cells expressing either SNED1^WT^ or SNED1^LAV^ for 48 hours. Interestingly, cells overexpressing SNED1^WT^ followed the disorganized alignment patterns of a SNED1^LAV^- containing ECM (**Figure 4E, *top right panel***), as quantified in **Figure 4F**. Vice-versa, cells overexpressing SNED1^LAV^ were able to align when seeded on a SNED1^WT^-containing ECM (**Figure 4E, *bottom left panel***), as quantified in **Figure 4F**. These results demonstrate that ECM architecture is sufficient to govern cell alignment and supersedes gene expression.

### Abrogation of the interaction of SNED1 with LDV integrins decreases cell proliferation

In addition to mediating cell organization, integrins transmit biochemical and mechanical signals to cells and initiate molecular cascades that govern various cellular processes, including cell proliferation (Jones and Jones, 2023). Indeed, we observed that cells expressing SNED1^LAV^ proliferated more slowly than cells expressing SNED1^WT^ (**Figure 4G**). We next sought to determine whether SNED1^WT^ or SNED1^LAV^-containing ECMs altered cell proliferation. To do so, we generated ECMs from SNED1^WT^- or SNED1^LAV^-expressing cells and decellularized them 9 days post-seeding. We then seeded *Sned1*^KO^ iMEFs on these decellularized ECMs. We used this cellular system to avoid interference from *Sned1* expression by re-seeded cells. 4 days post-reseeding, we found that cells seeded on a SNED1^LAV^-containing ECM had proliferated significantly less compared to cells seeded on a SNED1^WT^-containing ECM (**Figure 4H**), suggesting that SNED1 acts in a non-cell autonomous manner.

Since we have shown that the expression of SNED1^LAV^ had a profound impact on the organization of the entire ECM meshwork (*see above*), we sought to pinpoint whether SNED1 could directly alter cell proliferation. For this, we seeded *Sned1*^KO^ iMEFs on a cell-culture surface coated with purified SNED1^WT^ or SNED1^LAV^ and, found that cells were able to grow at the same rate on either substrate (**Figure 4I**). These data indicate that, rather than SNED1 directly signaling to cells via LDV-integrins, it is the consequences of the presence of SNED1^LAV^ on the entire ECM meshwork, including altered ECM alignment, and possible alterations in ECM mechanical properties, that modulate cell proliferation. This observation aligns with the established paradigm of mechanotransduction where ECM organization and stiffness govern cell proliferation, via the mechanosensitive YAP-TAZ pathway (Schwartz and Assoian, 2001; Sun et al., 2016; Totaro et al., 2018). In the future, it would be interesting to determine the impact of SNED1^LAV^ on the mechanical properties of the ECM and then delineate more precisely which SNED1-dependent signaling pathways are at play.

## CONCLUSION

The data presented in this short report provide the first evidence for a role between the novel ECM protein, SNED1, and LDV integrins in the assembly and organization of SNED1 fibrils. They also uncovered a role for SNED1 in the overall organization of the ECM meshwork, including of the meshwork composed of the two most abundant fibrillar ECM proteins, fibronectin and collagen I and in cell phenotypes, cell alignment and proliferation, both key to the pathophysiological processes we have previously incriminated SNED1 in, namely the migration of neural crest cells during developmental craniofacial morphogenesis (Barqué et al., 2021) and breast cancer metastasis (Naba et al., 2014). Our work thus paves the way to future studies aimed at determining to what extent SNED1/integrin interactions and SNED1-dependent ECM remodeling contribute to the *in-vivo* roles of SNED1.

## MATERIALS AND METHODS

### Plasmid constructs

The cDNA encoding full-length human SNED1 cloned into pCMV-XL5 was obtained from Origene (clone SC315884). Using a protocol previously described (Pally et al., 2025), we performed site-directed mutagenesis (Agilent, #200519) to introduce mutations to disrupt the two integrin binding sites RGD and LDV, alone (c.C120A; p.D40E; c.A932C; p.D311A) or in combination (c.C120A; p.D40E/c.A932C; p.D311A), in SNED1.

These constructs were then amplified by PCR to insert a FLAG tag at the C-terminus of SNED1 and the BglII and HpaI restriction sites to insert the constructs into the bicistronic retroviral vector pMSCV-IRES-Hygromycin. SNED1^WT^-FLAG was used for experiment testing the ability of cells to assemble exogenous SNED1, while SNED1LAV-FLAG was used to determine the ability of SNED1 to regulate cell proliferation.

The cloning of 6X-His-tagged versions of wild-type and mutant forms of SNED1 was previously described (Pally et al., 2025). His-tagged forms of SNED1 were used to coat cell culture dishes and determine the ability of SNED1 to mediate cell adhesion.

The cloning of GFP-tagged SNED1 (hereafter referred to as SNED1^WT^-GFP) was previously described (Vallet et al., 2021). Using site-directed mutagenesis (Agilent, #200519), we generated the following constructs: SNED1^RGE^-GFP (c.C120A; p.D40E), SNED1^LAV^-GFP (c.A932C; p.D311A), and SNED1^RGE/LAV^-GFP (c.C120A; p.D40E/c.A932C; p.D311A). These constructs were then shuttled into the bicistronic retroviral vector pMSCV-IRES-Puromycin between the BglII and EcoRI sites, as previously described (Vallet et al., 2021) and were used for all imaging experiments.

The sequences of the primers used for site-directed mutagenesis and subcloning are listed in **Supplemental Table S1**. All constructs were verified by Sanger sequencing at the UIC Genome Research Core facility.

### Cell culture

#### Cell maintenance

Immortalized mouse embryonic fibroblasts isolated from *Sned1*^KO^ mice (refer to as *Sned1*^KO^ iMEFs in the manuscript) (Barqué et al., 2021) stably overexpressing GFP-tagged wild type or integrin-binding mutant forms of SNED1 and human embryonic kidney 293T cells (further termed 293T) were cultured in Dulbecco’s Modified Eagle’s medium (DMEM; Corning, #10-017-CV) supplemented with 10% fetal bovine serum (FBS; Sigma, #F0926) and 2 mM glutamine (Corning, #25-005-CI), this medium will be further referred to as complete medium. All cell lines were maintained at 37 °C in a 5% CO2 humidified incubator.

#### Retrovirus production

293T cells were seeded in a 6-well plate at ∼30% confluency. The following day, cells were transfected using Lipofectamine 3000 (Invitrogen, #L3000008) with a mixture containing 1 µg of retroviral vector with the construct of interest and 0.5 µg each of a packaging vector (pCL- Gag/Pol) and vector encoding VSVG coat protein prepared in Opti-MEM (Gibco, #31985070). After 24 hours, the transfection mixture was replaced with fresh complete medium and cells were cultured for an additional 24 hours. The conditioned medium containing viral particles was collected, passed through a 0.45 µm filter, and either immediately used for cell transduction or stored at -80 °C for future use.

#### Generation of cells stably expressing different point mutants of SNED1

*Sned1^KO^* iMEFs were seeded at ∼30% confluency in a 6-well plate. The following day, cells were transduced with undiluted viral particles-containing conditioned medium. 24 hours after transduction, the medium was replaced with fresh complete medium, and cells were allowed to grow for an additional 24 hours before selection with puromycin (5 µg/ml). Once stable cell lines were established, the production and secretion of GFP-tagged mutant forms of SNED1 were confirmed using immunoblotting of total protein extracts (TE) and conditioned medium (CM), respectively, with an anti-SNED1 antibody; *see Results*. See **Supplemental Table S2** for details on antibodies used.

#### Preparation of fibronectin-depleted FBS

Fibronectin was depleted from FBS using a gelatin Sepharose 4B affinity resin (Cytiva, #17095601), as previously described (Sabatier et al., 2013; Sechler et al., 1996). All steps were performed on ice or in a cold room using ice-cold reagents. Gelatin Sepharose resin was supplied in 20% ethanol and was washed with 1X PBS before packing. 100 mL resin was packed into an Econo column using an Econo gradient pump (Biorad) with a maximum flow rate of 6.0 mL/min. Once packed, the column was equilibrated with 5 column volumes of 1X PBS at a flow rate of 4.5 mL/min. Following equilibration, 300 mL of FBS was loaded onto the column at a flow rate of 4.0 mL/min using an Econo pump and looped for 200 minutes. The flow-through was sterilized using a 0.2µm filter. Fibronectin-depleted FBS was used in the experiments to test the ability to assemble exogenous fibronectin or SNED1 into fibrillar structures.

### Protein purification

#### Conditioned medium collection

Conditioned medium containing recombinant SNED1 (WT or integrin-binding mutants) was collected as described previously (Pally et al., 2025). In brief, 293T cells stably expressing the protein of interest were seeded in a 15-cm dish in complete medium and allowed to grow until they reached 100% confluency. The culture medium was aspirated, and the cells were rinsed with 1X D-PBS containing Ca^2+^ and Mg^2+^ (D-PBS^++^, Cytiva, #SH30264.01). The medium was replaced with serum-free DMEM supplemented with 2 mM glutamine. Serum-free conditioned medium (SFCM) containing the secreted protein of interest was harvested after 48 hours, and cells were allowed to recover for 48 h in complete medium before repeating the next cycle. Cells were discarded after four to five cycles. At each collection time point, an EDTA-free protease inhibitor cocktail (0.067X final concentration; Thermo Fisher Scientific, #A32955) was added to the CM, and the CM was centrifuged at 2,576 × *g* for 10 minutes at 4 °C to remove any cells or cellular debris. Pre-cleared supernatants were collected and stored at −80 °C until further processing.

#### Purification of His-tagged and FLAG-tagged SNED1 proteins

Metal affinity protein purification of SNED1^WT^-His, SNED1^RGE^-His, SNED1^LAV^-His, and SNED1^RGE/LAV^-His was performed at the UIC Biophysics Core facility as previously described (Pally et al., 2025). Immunoprecipitation of FLAG-tagged forms of SNED1 (SNED1^WT^-FLAG, SNED1^RGE^-FLAG, SNED1^LAV^-FLAG, and SNED1^RGE/LAV^-FLAG) was performed in-house using an anti-FLAG resin (Sigma, #A220), as previously described (Pally et al., 2025).

### Immunoprecipitation of SNED1-GFP from conditioned medium

iMEFs overexpressing SNED1^WT^-GFP, SNED1^RGE^-GFP, SNED1^LAV^-GFP, or SNED1^RGE/LAV^-GFP were seeded at 5.5 x 10^5^ in a sterile tissue-culture-treated 6-well plate. When cells had reached 100% confluency (∼48h post-seeding), the culture medium was replaced and supplemented with 50 μg/mL ascorbic acid (Thermo Scientific, #352685000). The conditional medium (CM) was collected 4 days post-seeding and subjected to immunoprecipitation using protein A/G agarose beads (Thermo Scientific, #20421) and either IgG isotype as a negative control (Invitrogen, #PI31903) or an anti-GFP antibody (Sigma, #G6539; **Supplemental Table S2**). A second negative control, omitting antibodies, was also included. Samples were incubated on a rotating wheel at 4 °C, overnight. Samples were then centrifuged at 8,200 x *g* at 4 °C for 30 sec. to pellet the beads and immunoprecipitated proteins. Beads were washed three times in cold 1X PBS. Following the final centrifugation, beads were dried and immunoprecipitated proteins were resuspended in 3X Laemmli buffer containing 100 mM DTT. Samples were boiled for 10 minutes at 95 °C and then analyzed via SDS-PAGE and western blot.

### Deoxycholate (DOC) solubility assay

The DOC solubility assay was performed as previously described (Pankov and Yamada, 2004; Wierzbicka-Patynowski et al., 2004). iMEFs overexpressing either wild type or integrin-binding mutants of SNED1-GFP were seeded at 5.5 × 10^5^ in 6-well sterile cell culture-treated plates. Approximately 48 hours after seeding, 50 μg/mL ascorbic acid was added to each well. Every 48 hours following that, half of the medium was replaced with complete medium containing 100 μg/mL ascorbic acid. After a specified time in culture (3 days, 6 days, or 9 days), the conditioned medium (CM), containing proteins secreted by the cells, was collected and centrifuged for 3 minutes at 800 x *g* to remove any cellular debris, and the DOC solubility assay was performed on the cell culture layer composed of cells and their ECM. In brief, cells were washed with cold 1X PBS. 1 mL of DOC lysis buffer (5% deoxycholate, 3M Tris-HCl pH 8.8, 500 mM EDTA), protease inhibitors with EDTA (Thermo Scientific, #A32953), 167 μg/mL DNase I (Sigma, #AMPD1-1KT) was added to each well and incubated at 4°C for 1 hour. Samples were then collected using a sterile scraper and transferred to clean 1.5 mL tubes. Samples were sheared using a 26G needle to reduce viscosity and then centrifuged for 15 minutes at 21,100 × *g* at 4 °C. The supernatant (DOC-soluble material) was collected in a clean tube and stored at -80 °C. The pellet, containing DOC-insoluble proteins, was stored at -80 °C. Protein quantification of the DOC-soluble fraction was performed via BCA assay (Thermo Scientific, #23225), and samples were prepared for SDS-PAGE.

In parallel, total protein extracts (TE) from cells seeded and treated under the same conditions were collected by lysing the cells in 3X Laemmli buffer (0.1875 M Tris-HCl, 6% SDS, 30% glycerol) containing 100 mM dithiothreitol. Samples were then sheared using a 26G needle to reduce viscosity and boiled for 10 minutes at 95 °C and stored at -80 °C.

### SDS-PAGE and western blot

Protein samples were separated by SDS-PAGE at constant current (20 mA for stacking gel, 25 mA for resolving). Proteins were then transferred to nitrocellulose membranes at a constant current of 100V for 3 hours at 4 °C. Membranes were stained with Ponceau stain for 10 minutes and imaged to confirm the transfer of proteins. Membranes were then blocked in 5% non-fat milk in PBS containing 0.1% Tween-20 (PBST) for 60 minutes at room temperature under constant shaking. Membranes were then incubated in the presence of primary antibodies (**Supplemental Table S2**) in 5% non-fat milk in PBST overnight with constant shaking at 4 °C. The following day, membranes were washed three times with PBST, followed by incubation with horseradish peroxidase (HRP)-conjugated secondary antibodies (**Supplemental Table S2**) in 5% non-fat milk in PBST for 60 minutes with constant shaking at room temperature. Membranes were again washed three times with PBST. The presence of proteins of interest was detected by chemiluminescence, using the Pierce ECL Western blotting substrate (Thermo Scientific #32109). Imaging was performed using a ChemiDoc MP imaging system (Bio-Rad).

### Generation of cell-derived ECMs

#### Cell seeding

Cell-derived ECMs from iMEFs overexpressing GFP alone or GFP-tagged forms of SNED1 were prepared under aseptic conditions following a protocol previously established by the laboratory of Dr. Jean Schwarzbauer (Harris et al., 2018) with minor modifications. Briefly, glass coverslips were inserted in non-cell-culture-treated sterile plastic plates and coated with autoclaved and sterile-filtered 0.2% gelatin in 1X D-PBS^++^ for 1 hour at 37 °C. The gelatin solution was aspirated, and coverslips were washed with 1X D-PBS^++^. The gelatin coating was crosslinked using 1% glutaraldehyde (Electron Microscope Sciences, #16220) for 30 minutes at room temperature. The glutaraldehyde was aspirated, and coverslips were incubated for 5 minutes with 1X PBS, for a total of three washes. To quench any remaining glutaraldehyde, 1 M ethanolamine (Thermo Scientific, #451762500) was added to each well and incubated for 30 minutes at room temperature. The ethanolamine was aspirated, and coverslips were again washed for 5 minutes with 1X PBS, three times. After the final wash, complete medium was added to each well, and coverslips were stored at 37 °C until cell seeding. 2.0 x 10^5^ cells per 18 mm coverslip or 1.0 x 10^5^ cells per 12 mm coverslip were seeded. When cells reached 100% confluency (approximately 48 hours after seeding), the medium was replaced and supplemented with 50 μg/mL ascorbic acid (Thermo Scientific, #352685000). Every 48 hours following, half of the medium was replaced with complete medium containing 100 μg/mL ascorbic acid.

#### Decellularization

After a specified time in culture (3 or 9 days), cell layers were decellularized. Cells were washed twice in 1X PBS, then washed three times using pre-warmed wash buffer 1 (100 mM Na2HPO4, 2mM MgCl2, 2 mM EGTA, pH 9.60). Cells were then incubated with pre-warmed lysis buffer (8 mM Na2HPO4, 3% IGEPAL CA-360, pH 9.60) for 15 minutes at 37 °C. After adding the lysis buffer, the plate was tilted to aid cell lysis. Tilting was repeated every 5 minutes, for a total of three times. The lysis buffer was replaced for a second and then a third incubation with fresh, pre-warmed lysis buffer, for 40 minutes at 37 °C. Samples were then incubated with pre-warmed wash buffer 2 (10 mM Na2HPO4, 300 mM KCl, pH 7.50) containing 0.5% deoxycholate (Thermo Scientific, #J6228822) for 3 minutes, then washed three times with wash buffer 2, followed by four washes in deionized water at room temperature. 1X PBS was added to coverslips, and the samples were kept protected from light in aluminum foil at 4 °C until further processing. After each step, coverslips were visualized using a light microscope to assess cell removal.

### Assembly of exogenous SNED1 by cells

The assay was performed as previously described (Vega et al., 2020). In brief, 1.0 x 10^5^ cells were seeded on sterile 12-mm coverslips in DMEM supplemented with 2 mM glutamine and 10% fibronectin-depleted FBS.The medium was replaced the next day, and cells were incubated in the presence of 10 µg/mL of human plasma fibronectin (Millipore Sigma, #FC010) or SNED1^WT^- FLAG, and cultured for an additional 48 h. Samples were then fixed using ice-cold methanol:acetone (1:1) at -20 °C for 15 minutes, stained, and imaged as described below.

### Adhesion assay on purified SNED1 substrate

#### Substrate coating

Adhesion assays were performed as previously described (Humphries, 1998; Pally et al., 2025). In brief, the wells of a 96-well plate were coated with 50 µl of purified human SNED1-His (10 µg/mL) for 3 hours at 37 °C. Fibronectin (5 µg/ml; Millipore, #FC010) was used as a positive control. The adsorbed protein was immobilized with 0.5% (v/v) glutaraldehyde for 15 minutes at room temperature. To prevent cell adhesion to plastic, wells were blocked using 10 mg/ml heat-denatured bovine serum albumin (BSA; Sigma, #A9576). Uncoated wells blocked with BSA (10 mg/ml) alone were used as a negative control.

#### Cell seeding and crystal violet assay

A single-cell suspension of *Sned1*^KO^ iMEFs containing 5 × 10^5^ cells/ml was prepared in complete medium as previously described (Pally et al., 2025). 25,000 cells were added to each well and allowed to adhere for 30 minutes at 37 °C. Loosely attached or non-adherent cells were removed by washing the wells with 100 µl of 1X D-PBS^++^ three times, and remaining adherent cells were fixed using 5% glutaraldehyde for 30 minutes at room temperature. Cells were stained with 0.1% crystal violet (Alfa Aesar, #B21932) in 200 mM 2-(N-morpholino) ethanesulfonic acid (MES), pH 6.0, for 60 minutes at room temperature. After washing off the excess unbound crystal violet, the dye was solubilized in 10% acetic acid. Absorbance values were measured at λ=570 nm using a Bio-Tek Synergy HT microplate reader. Cell adhesion was determined by interpolating the absorbance values from a standard curve generated by seeding cells at several dilutions (10–100%) on wells coated with 0.01% poly-L-lysine (Sigma, #P4707) and fixing them directly by adding 5% (v/v) glutaraldehyde.

### Cell-ECM alignment

To assess the ability of cells to align on decellularized ECMs, 2.0 × 10^5^ cells were seeded onto decellularized ECMs produced by *Sned1*^KO^ iMEFs overexpressing SNED1^WT^-GFP or SNED1^LAV^-GFP after 9 days, and allowed to adhere and grow in complete medium for 48 h. Cells were fixed using 4% paraformaldehyde (PFA) (Electron Microscope Sciences, #15710), stained and imaged as described below.

### Cell adhesion and cell spreading on decellularized cell-derived ECMs

Decellularized ECMs produced by *Sned1*^KO^ iMEFs overexpressing SNED1^WT^-GFP or SNED1^LAV^- GFP were prepared as described above. 20,000 *Sned1*^KO^ iMEFs overexpressing GFP alone were seeded onto each coverslip and allowed to attach for 30 min (adhesion assay) or 3 hours (spreading assay) at 37 °C, 5% CO2 in a humidified chamber. Non-adherent cells were aspirated, and coverslips were washed with PBS, three times, to remove loosely attached cells. Adherent cells were fixed using 4% PFA and processed as described below. Coverslips were imaged using a Zeiss Axio Imager Z2 microscope with a 20X objective (adhesion assay) or a 40X objective (spreading assay). Cell counting was performed using ImageJ.

### Cell proliferation assay

Cell proliferation was quantified using the Methyl Thiazolyl Tetrazolium (MTT) assay as per the manufacturer’s protocol (Abcam, #ab211091). 5 × 10^3^ cells were seeded per well (uncoated or coated with either SNED1^WT^-FLAG or SNED1^LAV^-FLAG) in a 96-well plate in complete medium. Alternatively, 25 × 10^4^ cells were seeded on decellularized ECMs in complete medium. Cells were supplemented with fresh complete medium every 48 h. To measure cell proliferation, the complete medium was replaced with serum-free medium containing MTT reagent at a 1:1 ratio and incubated for 30 minutes at 37 °C in a 5% CO2 humidified incubator in the dark. The medium was aspirated and the formazan crystals were dissolved in MTT solvent at room temperature for 15 minutes on an orbital shaker. The optical density (OD) was measured at λ=590 nm using a Bio- Tek Synergy HT microplate reader. The MTT solvent alone was used as a blank.

### Immunofluorescence staining

Unless stated otherwise, cells or decellularized ECMs were fixed with 4% PFA for 15 minutes at room temperature. Free aldehyde groups were quenched using 50 mM NH4Cl for 15 min at room temperature. Following fixation, samples are rinsed three times with 1X D-PBS^++^ and stained immediately or stored in 1X PBS at 4 °C for future use.

#### Staining of decellularized ECMs

Decellularized and fixed ECMs were stored for 96 hours at 4 °C in 1X PBS. Samples were blocked with 1% bovine serum albumin (BSA) in 1X PBS containing 0.1% Triton X-100 at room temperature for 1 hour to prevent non-specific binding. Samples were then incubated with primary antibodies (**see Supplemental Table S2**) in 1X PBS containing 0.05% Triton X-100 overnight at 4 °C. The following day, coverslips were washed three times with 1X D-PBS^++^ containing 0.1% Triton X-100 and then incubated with secondary antibodies in 1X PBS + 0.05% Triton X-100 for 1 hour at room temperature. Coverslips were washed three times with 1X D-PBS^++^ + 0.1% Triton X-100, then mounted on glass microscope slides using Fluoromount G. The slides were stored at 4 °C overnight. Coverslips were imaged using either the Zeiss Axio Imager Z2 or the Zeiss Confocal LSM 880, as indicated in the figure legends.

#### Staining of the actin cytoskeleton

Fixed cells were permeabilized using 0.25% Triton X-100 in 1X PBS for 10 minutes at room temperature and stained with phalloidin-AlexaFluor 647 (Invitrogen, #A22287) and 4′,6- diamidino-2-phenylindole (DAPI; Invitrogen #D1306) for 1 hour at room temperature. Coverslips were washed three times with 1X D-PBS^++^ containing 0.1% Triton X-100 for 5 minutes at room temperature. Coverslips were mounted on glass slides using Fluoromount G (Invitrogen, #00- 4958-02) and allowed to dry overnight at 4 °C before imaging.

### Imaging

#### Image acquisition

Stained samples were imaged using a Zeiss Axio Imager Z2 epifluorescence microscope or a Zeiss Confocal LSM 800 microscope, as indicated.

#### Image analysis

##### ECM thickness analysis

Z-stacks obtained via confocal microscopy were analyzed to measure the thickness of different ECM layers. The bottom-most (beginning) and top-most (end) slices where a non-noise (*i.e.*, specific) signal was observed were recorded. Thickness was calculated by taking the total thickness of the Z-stack divided by the total number of slices to obtain the thickness of each slice. The beginning slice number was subtracted from the end slice number, and the resulting value was multiplied by the slice thickness (0.33-0.35 μm) to obtain the protein layer thickness value. These values were calculated for 3 to 5 fields per coverslip, for each protein.

#### 3D reconstruction

The volume of each Z-stack was visualized using Fiji ImageJ. Volume Viewer plugin was used to analyze stacks, with the Z-aspect set to 10.0 in Max Projection mode with Tricubic smooth interpolation. XY, XZ, and XY snapshots were obtained for each field.

#### Colocalization analysis

Colocalization analysis was performed on Z-stacks using Zeiss ZEN. Thresholds were set using negative controls, and analyses between channels for each z-slice were recorded using Microsoft Excel. Line graphs using the Manders correlation coefficient were then plotted to visualize the changes across the specified portion of each Z-stack.

#### Area fraction density analysis

ECM density/area fraction was measured using Fiji ImageJ. 8-bit orthogonal projection (maximum intensity) images were imported and brightness and contrast were adjusted manually. Area fraction was selected in the analyze > set measurements table to obtain a staining density value for 3 to 5 fields per coverslip.

#### Fiber alignment analysis

The OrientationJ plugin was used to analyze F-actin alignment in cells and alignment of ECM fibers (Püspöki et al., 2016; Rezakhaniha et al., 2012). In brief, 8-bit orthogonal maximum intensity projection images were imported for measurement. The images were aligned along the dominant horizontal axis using the horizontal alignment tool in OrientationJ plugin. a 1-pixel local window was specified, and the cubic spline gradient was moved across the image to calculate structure tensors to determine local orientations. Orientation in degrees, ranging from 90° to +90° from the horizontal, was obtained as a histogram of the distribution of orientations.

### Statistical analysis

All experiments were performed at least three biological replicates unless mentioned specifically in respective figure legends. Data is represented as individual experimental values, mean or mean ± standard deviation, as indicated in the respective figure legends. Unpaired *t*-tests with Welch’s correction was performed to assess statistical significance in all analyses. Graphs were generated using GraphPad Prism.

## ACKNOWLEDGEMENTS

The authors would like to thank all the members of the Naba lab for insightful discussions and Pr. Martin Humphries (University of Manchester) for early feedback on this project.

## COMPETING INTERESTS

The Naba laboratory holds a sponsored research agreement with Boehringer-Ingelheim for work not related to the content of this manuscript.

## FUNDING

This work was supported by the National Institute of General Medical Sciences of the National Institutes of Health to AN [R01GM148423]. LP was supported by a T32 fellowship from the Vascular Biology, Signaling and Therapeutics training program [HL144459] and a UIC Graduate College - Dean’s Scholar Fellowship. AJ was supported by awards from the UIC Liberal Arts and Sciences Undergraduate Research Initiative (LASURI) and a UIC Honors College Undergraduate Research Grant.

## DATA AVAILABILITY

All relevant data can be found within the article and its Supplemental information. Research materials are available upon request via email to Dr. Naba.

## SUPPLEMENTAL FIGURE LEGENDS

**Supplemental Figure S1.**
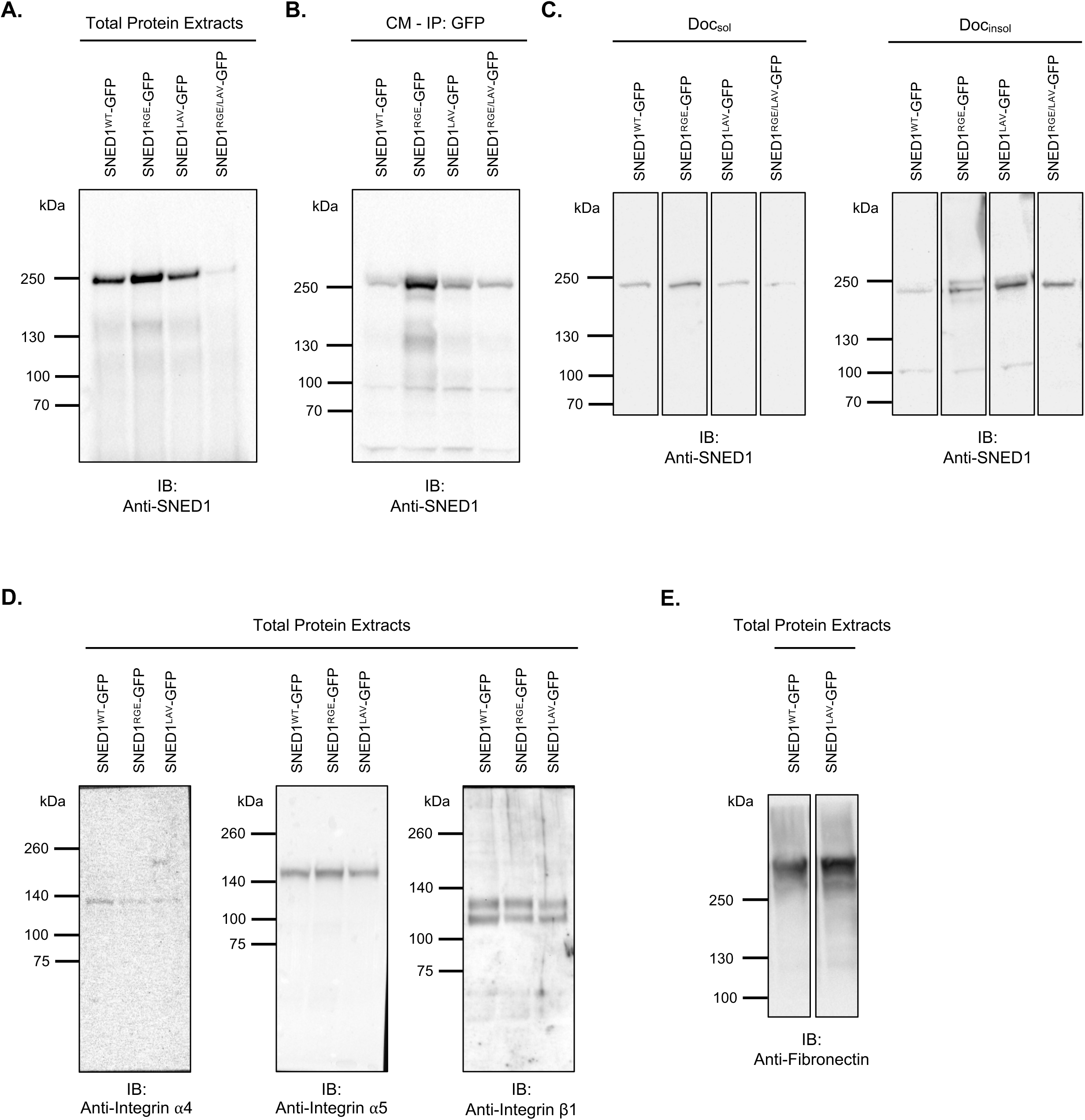
Expression of SNED1 and integrins in *Sned1*^KO^ iMEFs overexpressing SNED1^WT^-GFP, SNED1^RGE^-GFP, or SNED1^LAV^-GFP. A. Full-size immunoblots show the presence of SNED1-GFP in total protein extracts collected from *Sned1*^KO^ iMEFs overexpressing SNED1^WT^-GFP, SNED1^RGE^-GFP, or SNED1^LAV^-GFP, 4 days post-seeding. Images are representative of three independent biological replicates. B. Immunoblots show the presence of immunoprecipitated SNED1-GFP in the culture medium conditioned (CM) by *Sned1*^KO^ iMEFs overexpressing SNED1^WT^-GFP, SNED1^RGE^-GFP, or SNED1^LAV^-GFP, 4 days post-seeding. Images are representative of three independent biological replicates. C. Immunoblots show the presence of SNED1-GFP in the DOC-soluble (DOCsol) and DOC- insoluble (DOCinsol) fractions collected from *Sned1*^KO^ iMEFs overexpressing SNED1^WT^-GFP, SNED1^RGE^-GFP, or SNED1^LAV^-GFP, 2 days post seeding. Images are representative of three independent biological replicates. D. Immunoblots show the presence of α4, α5, and β1 integrins in total protein extracts from *Sned1^KO^* iMEFs overexpressing SNED1^WT^-GFP, SNED1^RGE^-GFP, or SNED1^LAV^-GFP. Images are representative of three independent biological replicates. E. Immunoblot shows fibronectin expression in total protein extracts from cells overexpressing SNED^WT^-GFP and SNED1^LAV^-GFP.

**Supplemental Figure S2.**
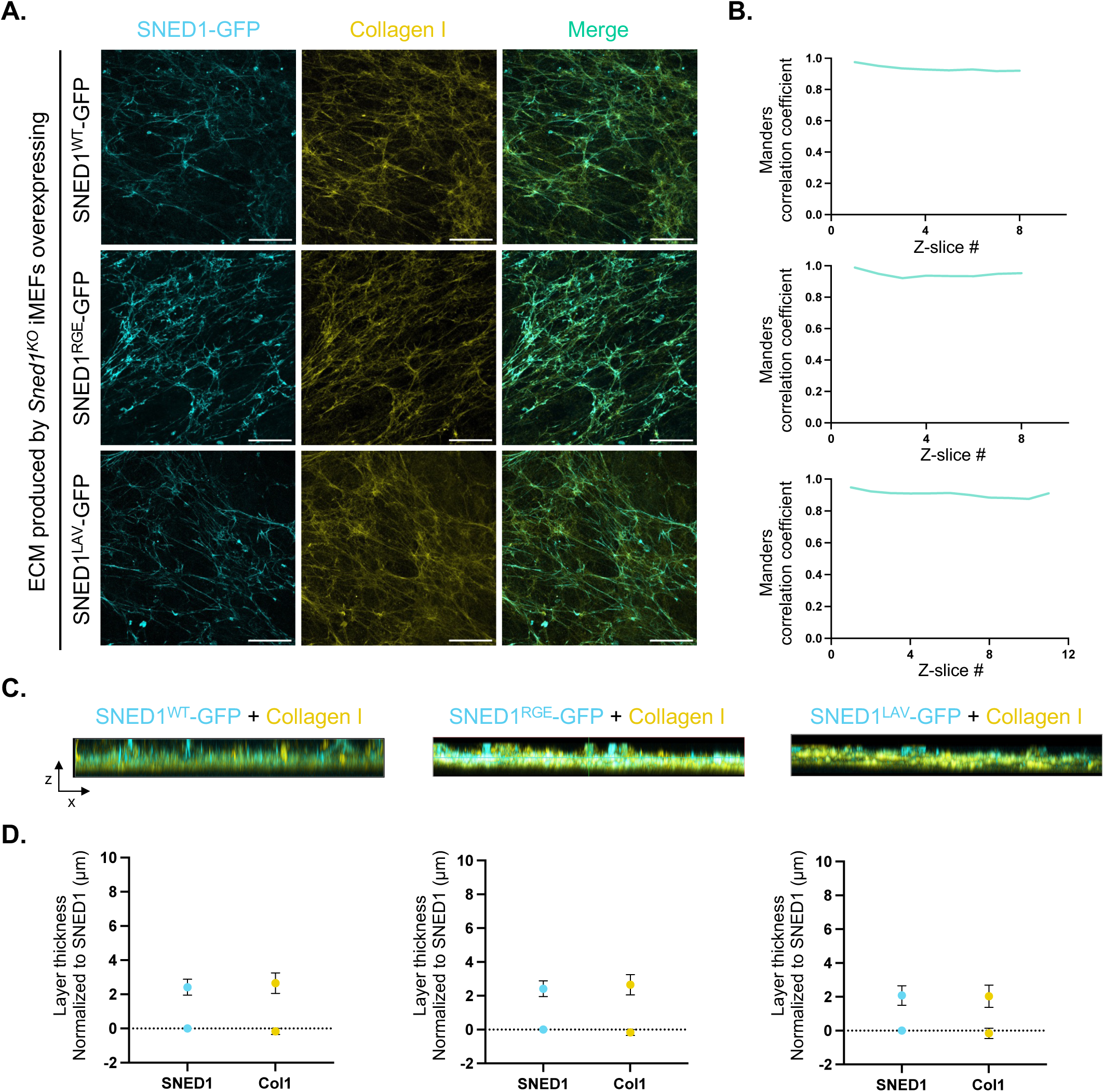
Disruption of integrin-binding motifs in SNED1 does not affect the colocalization of SNED1 with collagen I at the early stages of ECM assembly. **A.** XY orthogonal maximum projections show SNED1 (cyan) and collagen (yellow) fibers in the ECM produced by *Sned1*^KO^ iMEFs overexpressing SNED1^WT^-GFP (*top panels*), SNED1^RGE^-GFP (*middle panels*), or SNED1^LAV^-GFP (*bottom panels*) and decellularized 3 days post-seeding. Merge panels show overlap between SNED1 and collagen I signals. Scale bar: 30 µm. Images are representative of at least three independent biological replicates. **B.** Representative line graphs show Manders overlap coefficient between SNED1 and collagen I for each Z-slice where in-focus signal was detected in the ECM produced by *Sned1*^KO^ iMEFs overexpressing SNED1^WT^-GFP (*top panel*), SNED1^RGE^-GFP (*middle panel*), or SNED1^LAV^-GFP (*bottom panel*), and decellularized 3 days post-seeding. **C.** XZ orthogonal maximum projections show collagen I and SNED1-GFP signal in the ECMs produced by *Sned1*^KO^ iMEFs overexpressing SNED1^WT^-GFP (*left panel*), SNED1^RGE^-GFP (*middle panel*), or SNED1^LAV^-GFP (*right panel*), and decellularized 3 days post-seeding. Images are representative of three independent biological replicates. **D.** Dot plots depict the overlap of SNED1 and collagen I (ColI) signals in ECMs produced by *Sned1*^KO^ iMEFs overexpressing SNED1^WT^-GFP (*left panel*), SNED1^RGE^-GFP (*middle panel*), or SNED1^LAV^-GFP (*right panel*), and decellularized 3 days post-seeding. Lower and higher data points represent the beginning and end of in-focus signal detected in the ECM. Data points are normalized to the Z-slice value where SNED1’s signal is detected. Data is represented as mean ± standard deviation. Unpaired *t*-test with Welch’s correction did not show any statistical significance.

**Supplemental Figure S3.**
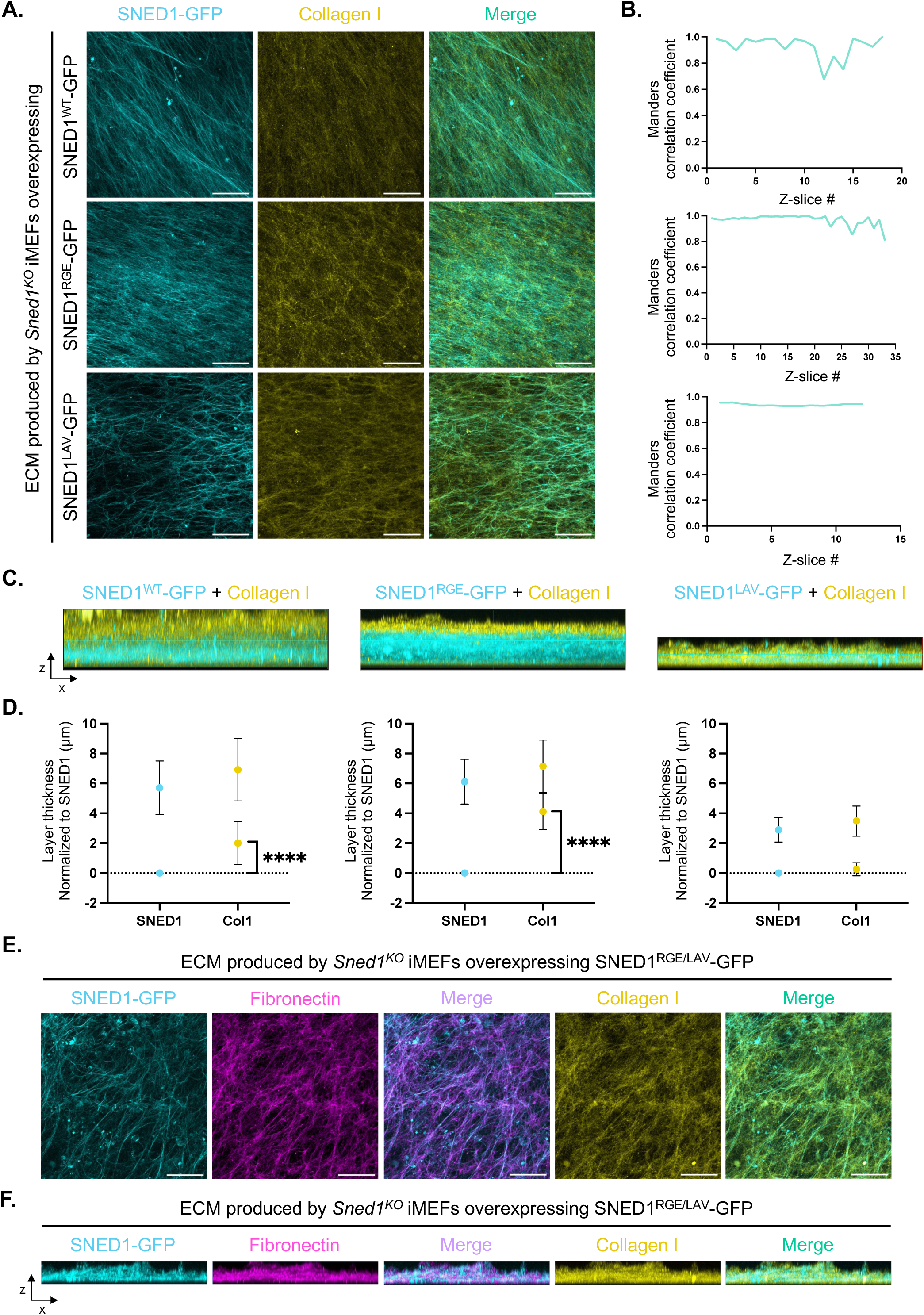
Disruption of the LDV but not the RGD integrin-binding motif in SNED1 impairs collagen I remodeling. **A.** XY orthogonal maximum projections show SNED1 (cyan) and collagen (yellow) fibers in the ECM produced by *Sned1*^KO^ iMEFs overexpressing SNED1^WT^-GFP (*top panels*), SNED1^RGE^-GFP (*middle panels*), or SNED1^LAV^-GFP (*bottom panels*), and decellularized 9 days post-seeding. Merge panels show overlap between SNED1 and collagen I signals. Scale bar: 30 µm. Images are representative of at least three independent biological replicates. **B.** Representative line graphs show Manders overlap coefficient between SNED1 and collagen I for each Z-slice where in-focus signal was detected in the ECM produced by *Sned1*^KO^ iMEFs overexpressing SNED1^WT^-GFP (*top panel*), SNED1^RGE^-GFP (*middle panel*), or SNED1^LAV^-GFP (*bottom panel*) and decellularized 9 days post-seeding. **C.** XZ orthogonal maximum projections show collagen I (yellow) and SNED1-GFP (cyan) signals in the ECMs produced by *Sned1*^KO^ immortalized mouse embryonic fibroblasts (*Sned1*^KO^ iMEFs) overexpressing SNED1^WT^-GFP (*left panel*), SNED1^RGE^-GFP (*middle panel*), or SNED1^LAV^-GFP (*right panel*), and decellularized at 9 days post cell-seeding. Images are representative of three independent biological replicates. **D.** Dot plots depict the overlap between SNED1 and collagen I (ColI) signals in ECMs produced by *Sned1*^KO^ iMEFs overexpressing SNED1^WT^-GFP (*left panel*), SNED1^RGE^-GFP (*middle panel*) or SNED1^LAV^-GFP (*right panel*), decellularized at 9 days post-seeding. Lower and higher data points represent the beginning and end of in-focus signal detected in the ECM. Data points are normalized to the Z-slice value where SNED1’s signal is detected. Data is represented as mean ± standard deviation. Unpaired *t*-test with Welch’s correction was performed to test statistical significance. ****p<0.0001. **E.** XY orthogonal maximum projections show SNED1 (cyan), fibronectin (magenta), and collagen (yellow) fibers in the ECM produced by *Sned1*^KO^ iMEFs overexpressing SNED1^RGE/LAV^-GFP and decellularized 9 days post-seeding. Merge panels show overlap between SNED1 and fibronectin or SNED1 and collagen I. Scale bar: 30 µm. Images are representative of one biological replicate. **F.** XZ orthogonal maximum projections show SNED1 (cyan), fibronectin (magenta), collagen (yellow) fibers in the ECM produced by *Sned1*^KO^ iMEFs overexpressing SNED1^RGE/LAV^-GFP and decellularized 9 days post-seeding. Merge panels show overlap between SNED1 and fibronectin or SNED1 and collagen I.

**Supplemental Figure S4.**
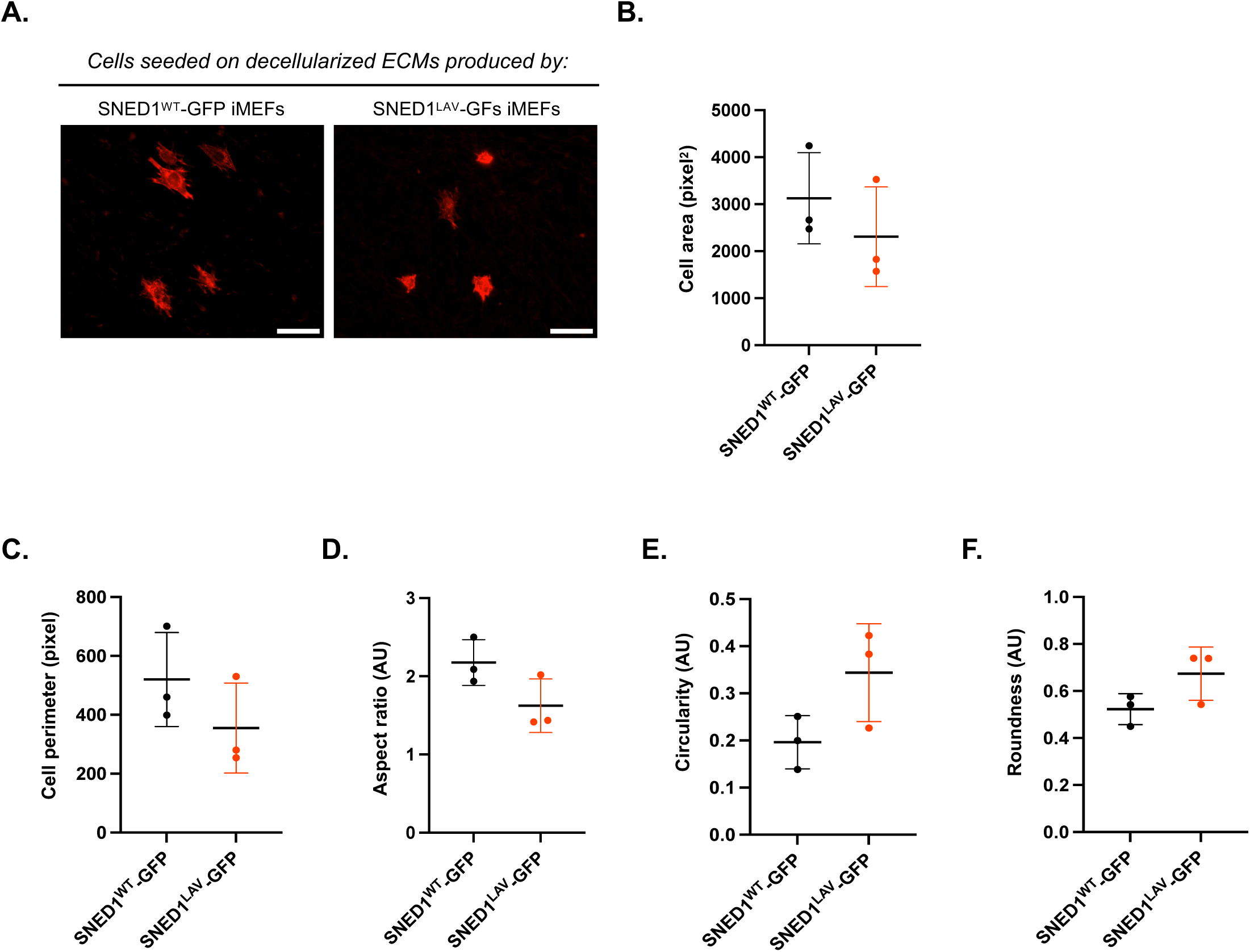
Mutation of the LDV integrin-binding motif in SNED1 moderately affects cell spreading. **A.** Microscopy images show the spreading of *Sned1*^KO^ iMEFs overexpressing GFP alone (visualized through phalloidin-AF647 staining) seeded on decellularized ECMs produced by *Sned1*^KO^ iMEFs overexpressing SNED1^WT^-GFP (*left panel*) or SNED1^LAV^-GFP (*right panel*) for 9 days. Images are representative of three biological replicates. Scale bar: 50 µm. **B.** Dot plot depicts the area of *Sned1*^KO^ iMEFs overexpressing GFP alone seeded on decellularized ECMs produced by *Sned1*^KO^ iMEFs overexpressing SNED1^WT^-GFP (black) or SNED1^LAV^-GFP (red) for 9 days. Data is represented as mean ± standard deviation from three biological replicates, with at least 10 imaging fields per coverslip, and at least two coverslips per replicate. Unpaired *t*- test with Welch’s correction did not show any statistical significance. **C-F.** Dot plots show a moderate decrease in cell perimeter (**C**), aspect ratio (**D**), and increased circularity (**E**) and roundness (**F**) of cells seeded on ECMs produced by *Sned1^KO^* iMEFs overexpressing SNED1^WT^-GFP (black) or SNED1^LAV^-GFP (red) and decellularized at 9 days post-seeding. Data is represented as mean ± standard deviation from three biological replicates, with at least 10 fields per coverslip and at least two coverslips per replicate. Unpaired *t*-test with Welch’s did not show any statistical significance.

## SUPPLEMENTAL TABLES

**Supplemental Table S1.**
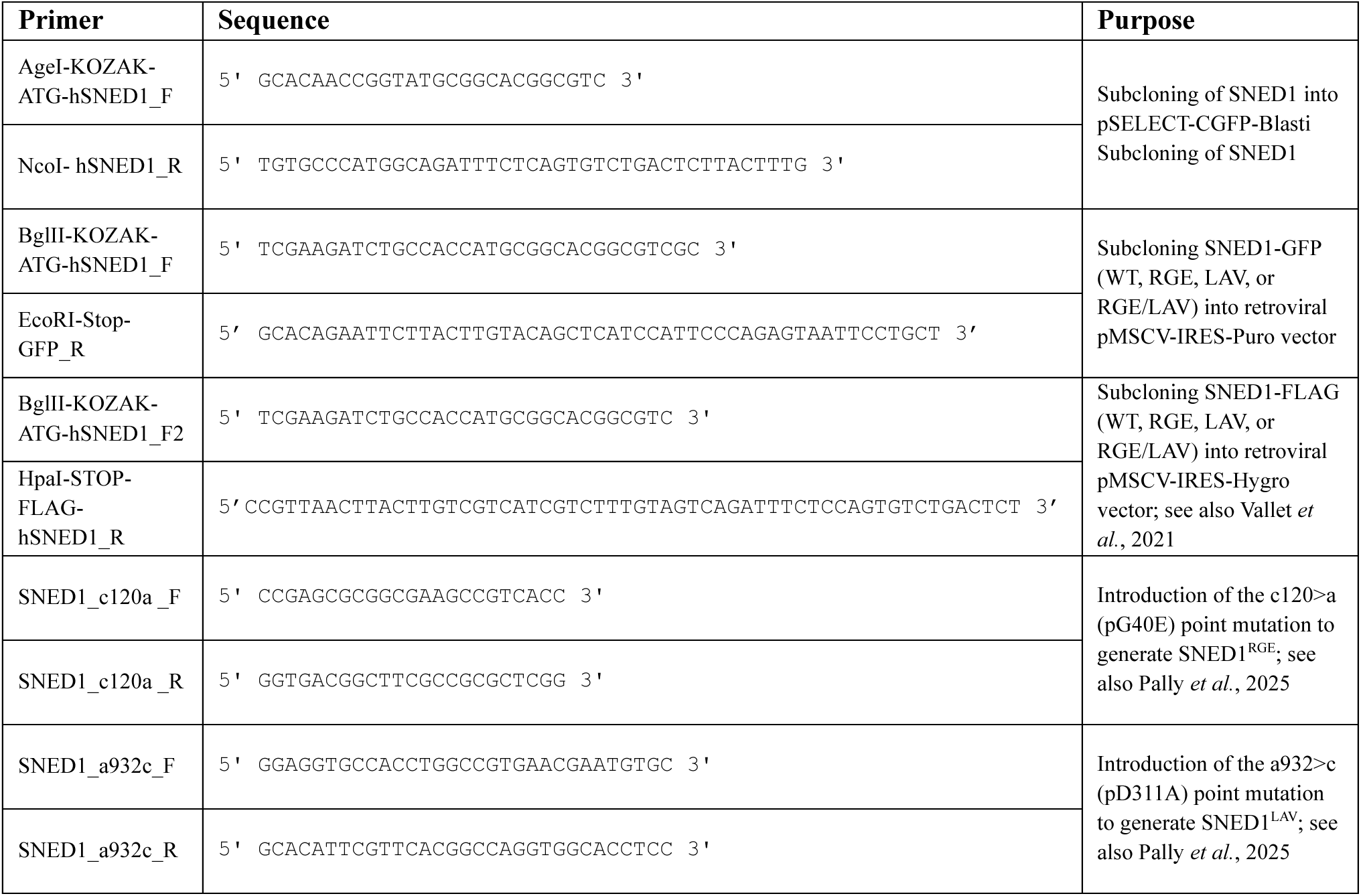
List of primers used in this study.

**Supplemental Table 2.**
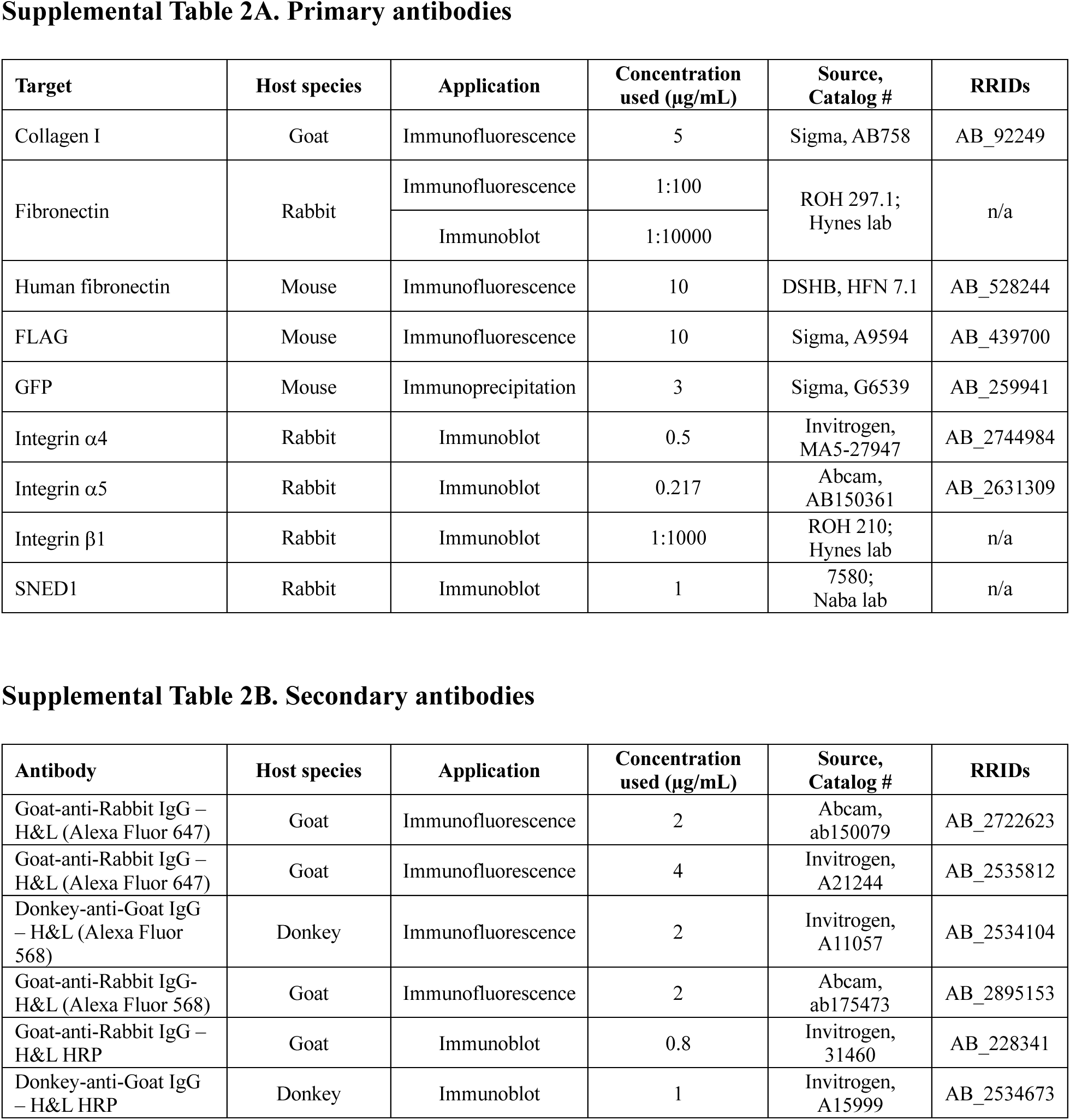
List of antibodies used for immunostaining and immunoblotting.

## Notes

### Competing Interest Statement

The authors have declared no competing interest.

